# Rare disruptive variants in the DISC1 Interactome and Regulome: association with cognitive ability and schizophrenia

**DOI:** 10.1101/083816

**Authors:** Shaolei Teng, Pippa A. Thomson, Shane E. McCarthy, Melissa Kramer, Stephanie Muller, Jayon Lihm, Stewart Morris, Dinesh Soares, William Hennah, Sarah Harris, Luiz Miguel Camargo, Vladislav Malkov, Andrew M McIntosh, J. Kirsty Millar, Douglas Blackwood, Kathryn L. Evans, Ian J. Deary, David J. Porteous, W. Richard McCombie

**Affiliations:** Stanley Institute for Cognitive Genomics, Cold Spring Harbor Laboratory, Cold Spring Harbor, NY, USA; Department of Biology, Howard University, Washington DC, USA; Centre for Genomic and Experimental Medicine, MRC/University of Edinburgh Institute of Genetics & Molecular Medicine, Western General Hospital, Edinburgh, UK; Centre for Cognitive Ageing and Cognitive Epidemiology, Edinburgh, UK; Institute for Molecular Medicine, Finland FIMM, University of Helsinki, Helsinki, Finland; UCB New Medicines, One Broadway, Cambridge, MA, USA; Genetics and Pharmacogenomics, MRL, Merck & Co, Boston, MA, USA; Division of Psychiatry, University of Edinburgh, Royal Edinburgh Hospital, Edinburgh, UK; Department of Psychology, University of Edinburgh, Edinburgh, UK.

## Abstract

Schizophrenia (SCZ), bipolar disorder (BD) and recurrent major depressive disorder (rMDD) are common psychiatric illnesses. All have been associated with lower cognitive ability, and show evidence of genetic overlap and substantial evidence of pleiotropy with cognitive function and neuroticism. *Disrupted in schizophrenia 1* (DISC1) protein directly interacts with a large set of proteins (DISC1 Interactome) that are involved in brain development and signaling. Modulation of DISC1 expression alters the expression of a circumscribed set of genes (DISC1 Regulome) that are also implicated in brain biology and disorder. Here, we report targeted sequencing of 59 DISC1 Interactome genes and 154 Regulome genes in 654 psychiatric patients and 889 cognitively-phenotyped control subjects, on whom we previously reported evidence for trait association from complete sequencing of the DISC1 locus. Burden analyses of rare and singleton variants predicted to be damaging were performed for psychiatric disorders, cognitive variables and personality traits. The DISC1 Interactome and Regulome showed differential association across the phenotypes tested. After family-wise error correction across all traits (FWER_*across*_), an increased burden of singleton disruptive variants in the Regulome was associated with SCZ (FWER_*across*_ *P*=0.0339). The burden of singleton disruptive variants in the DISC1 Interactome was associated with low cognitive ability at age 11 (FWER_*across*_ *P*=0.0043). These results suggest that variants in the DISC1 Interactome effect the risk of psychiatric illness through altered expression of schizophrenia-associated genes. The biological impact of rare variants highlighted here merit further study.

## INTRODUCTION

Schizophrenia (SCZ), bipolar disorder (BD) and recurrent major depressive disorder (rMDD) affect tens of millions of people worldwide. These disorders are moderately heritable and family history is a strong predictor of risk. Genome wide association studies (GWAS), structural variant analyses, and genome sequencing studies have identified that common single nucleotide variants (SNVs), low penetrant rare SNVs, moderate to high penetrant copy number variants (CNVs), and potentially causal *de novo* mutations each play a role in the genetic etiology of SCZ and BD and, to a lesser extent, in rMDD.^1–4^

There is now strong evidence for shared genetic risk across traditional diagnostic boundaries supporting the observation of ‘mixed’ diagnoses families.^5,6^ GWAS studies capture common, ancient variation and point to an additive, polygenic architecture that transcends psychiatric diagnoses to predict cognitive ability variables.^1,7,8^ Lower cognitive function, both premorbid and post-onset, has been associated with these disorders, and recently polygenic risk score analysis has suggested a small, but significant genetic correlation between risk for major mental illness and cognitive ability.^9^

In a complementary fashion to common variants, rare variants have been identified that segregate with psychiatric disorder in a quasi-Mendelian fashion and impact upon normal cognitive function.^10,11^ One such example is a balanced t(1;11) (q42;q14) translocation in *Disrupted in Schizophrenia 1* (*DISC1*) gene which was first identified at the breakpoint in a large Scottish pedigree highly burdened with SCZ, BD and rMDD.^12^ Independent reports of linkage and association have since reported evidence for region-wide association of *DISC1* variants, or more commonly haplotypes, with these and other psychiatric disorders as well as for cognitive and neuropsychological traits^13–15^. However, *DISC1* is not a GWAS significant finding.^16,17^ Recently, we reported deep sequencing of the *DISC1* locus (528kb) in 1 542 samples that identified 2 010 rare variants, of which ~60% were novel.^17^ We identified a common intronic variant with region-wide association for rMDD and also identified a rare missense mutation (R37W), previously reported in a SCZ case^18^, in an individual with rMDD and in additional family members with mental disorders.^17^ R37W has now been shown to have a direct effect on mitochondrial trafficking.^19,20^ Burden analysis also identified nominal associations with measures of depressed mood and cognitive ability at age 11, age 70 and cognitive ageing (change in cognitive ability between age 11 and age 70). Motivated by these findings, we hypothesized that further insights might emerge from a directly comparable study of the DISC1 pathway genes.^21–24^

Molecular studies have shown that DISC1 functions as a scaffold protein that is critical in cell signaling, neuronal development and ontogenesis through multiple protein-protein interactions.^25^ DISC1 interacting partners (DISC1 Interactome) are enriched for proteins known to be involved in neural proliferation, migration, signaling and synaptic function.^14,21,22,24,26^ Positive case-control associations have been reported with psychiatric disorders for the following DISC1 Interactome genes: *ATF4, CIT, NDE1, PCM1, PDE4B, PDE4D* and *YWHAE*.^14^ Structural rearrangements of *PDE4B* and *NDE1* have been reported in patients with SCZ.^27,28^ Sequencing studies in SCZ have examined the burden of variants in *DISC1* and its interacting partners with mixed results. Moens *et al.* Sequenced *DISC1* and ten interacting genes in 486 SCZ cases and 514 controls and observed an excess of rare missense variants in affected individuals.^29^ Kenny *et al.* sequenced the exons of 215 synaptic genes including *DISC1* and 22 interaction partners in 147 individuals with autism, 273 with SCZ and 287 controls.^30^ There was no enrichment of loss of function (LoF) mutations in the subset of DISC1 interacting genes, but singleton LoF variants were identified in *DISC1*, *DMD* and *TRAF3IP1* in patients with autism. Although recent exome based studies of SCZ did not analyze DISC1 interacting genes directly, an over representation of *de novo* and rare variation was observed in genes sets such as the activity-regulated cytoskeleton-associated scaffold protein complex at the postsynaptic density.^31^ This finding suggests that the DISC1 Interactome may be enriched in genetic risk factors that function through perturbing neuronal development and function.^31^

In addition to the DISC1 Interactome, disruption of the expression of *DISC1*, or its interactors, has been shown to alter the expression of a further set of genes (DISC1 Regulome).^23,32,33^ This gene set is enriched for neurodevelopmental, synaptogenic and sensory perception genes as well as for registered drug targets for psychiatric and neurological disorders.^23,26^ We selected DISC1 Regulome genes on the basis of prior evidence of genetic association with disruption of *DISC1*, or for haplotype or model dependent expression patterns correlated with *DISC1* expression,^34^ *DISC1* alleles,^23^ or with DISC1 Interactome genes.

Here, we report the targeted sequencing of 59 DISC1 Interactome genes (including *TSNAX* and *DISC1*) and 154 DISC1 Regulome genes. We report rare variants with a minor allele frequency (MAF) less than 1% and singletons across the exons of these gene sets in the same cohort of subjects in which the full DISC1 locus (528kb) was sequenced and reported.^17^ Of the 1 543 subjects, 1 446 samples passed quality control, comprising 575 cases with SCZ, BD or rMDD, and 871 generally healthy older people who acted as controls. As for our previous study, we report the gene-wide and gene-set burden analysis of rare variants and singletons with respect to psychiatric disorders, associated personality traits, and cognitive variables. We focused our analysis on the accumulation rate and burden of rare variants (MAF<1%) and the subset of singleton variants stratified into three damaging mutation categories: (1) disruptive mutations, (2) non-synonymous strictly damaging mutations (*NS*_*strict*_), and (3) non-synonymous broadly damaging mutations (*NS*_*broad*_).

## MATERIALS AND METHODS

The materials and methods are described in full in the Supplementary Information. Briefly, we analyzed 1543 DNA samples comprising 654 cases (241 SCZ, 221 BD and 192 rMDD) from Scottish hospital patients and 889 community-dwelling, generally healthy older people from the Lothian Birth Cohort of 1936 (LBC1936), as described previously.^17^

A total of 213 genes were selected for sequence analysis (Supplementary Table S1). The DISC1 locus (*DISC1*, *TSNAX* and *TSNAX-DISC1*) and 56 direct DISC1 protein-protein interactors defined the DISC1 Interactome gene set. 154 additional genes related to DISC1 expression from previous microarray analyzes comprised the DISC1 Regulome gene set. Genomic regions comprising approximately 11.7Mb (3.3Mb exons) were captured using a custom solution capture probe set (Roche NimbleGen). Each sample capture was sequenced using a HiSeq2000 sequencer (Illumina). Sequence reads were aligned to the human NCBI Build 36 (hg18) reference using BWA.^35^ Variant calling was performed using GATK,^36^ and high quality SNVs were filtered by standardized filtering parameters. Using PLINK,^37^ we applied data quality control filters as described previously^17,38^ to exclude samples and SNVs that introduce bias (Supplementary Figure S2-S5). Sanger sequencing was used to optimize the quality control filters and exclude all identified false positive SNVs from further analysis. SNVs were matched to hg19 coordinates using liftOver from UCSC, and ANNOVAR^39^ was used for variant annotation based on the human reference genome hg19 (RefSeq). The coding variants were grouped into three mutation classes, similar to previous analyses,^4,31^ based on predicted functional effects: Disruptive, nonsense and splice site variants; NS_*strict*_, disruptive plus missense variants predicted as damaging by all five algorithms (SIFT,^40^ PolyPhen2 HumDiv and HumVar,^41^ LRT^42^ and MutationTaster^43^); NS_*broad*_, disruptive plus missense variants predicted as damaging by at least one of the algorithms above. The burden and accumulation rate of rare variants (MAF<1%) and singletons in these mutation classes was assessed in each of the case cohorts and a combined cohort of all diagnoses; and for cognitive measures and personality traits. The gene set and gene-wide burden analyses for all genes containing more than one rare variant were performed using the R package ‘SKAT’.^44^ Exact Poisson tests were performed in R to evaluate the accumulation rates of singletons and rare variants under a Poisson distribution in cases compared to controls in each functional mutation class. A bootstrap resampling approach (n=10 000) was used to estimate the significance of all tests (Family-Wise Error Rate, FWER), taking into account multiple tests within each diagnosis (FWER_within_) and across all diagnoses (FWER_across_).

## RESULTS

### Targeted Sequencing and Genetic Discovery in 213 DISC1 Interactome and Regulome Genes

We sequenced the exons (3.3Mb), promoters, and conserved regions (a total of 11.7Mb) of 213 genes functionally related to DISC1 in a cohort of 1 543 Scottish individuals. Of these, 1 464 samples (95%) were sequenced to a minimum coverage depth of 20x across 80% of the targeted bases (Supplementary Table S2). Coverage was uniform across all sample groups (Supplementary Figure S1). Following sequence- and variant-based quality filters, 196 080 SNVs in 1 446 samples (211 cases of SCZ, 169 cases of rMDD, 195 cases of BD, and 871 controls from the LBC1936) were left for further analyses (Supplementary Table S2, Figure S2-S5). Of the 196 080 SNVs, 78% have a MAF less than 1%. Only 40% are reported in the 1000 Genome Project European subset (Supplementary Table S3). Based on RefSeq functional annotations using ANNOVAR, 169 905 SNVs mapped to introns, 5 410 to 5’/3’ UTRs, and 4 523 to coding regions. Of the 4 523 exonic variants, 1 893 were functionally classified with respect to coding potential as silent variants, 2 569 as missense, and 41 as nonsense. A further 24 SNVs were annotated as splice site variants. SNVs showing greater functional impacts on protein function are more likely to be rare: 100% of nonsense and 92% of splice site variants have MAF less than 1%, compared to 79% of silent and 78% of intronic variants.

### Analysis of Genetic Variation in the DISC1 Interactome and Regulome with Psychiatric Illness

#### Rare Functional Variant Analysis in the DISC1 Interactome

There was no significant burden of rare disruptive, NS_*strict*_ or NS_*broad*_ variants in SCZ, BD, or rMDD nor in a combined cohort of all diagnoses compared to controls in the DISC1 Interactome (Supplementary Table S6). There was a nominal association of fewer rare disruptive variants in SCZ (unadjusted *P*=0.0188), but no significant difference between the accumulation rate of rare variants for any diagnosis after Family-Wise Error Rate (FWER) correction (Supplementary Table S7). None of the proportions of NS_*strict*_ and NS_*broad*_ rare or singleton variants deviated from the null hypothesis after FWER_across_ correction.

The gene-wide burden of non-synonymous coding changes was nominally, but significantly increased in psychiatric disorders (*P*=0.0048-0.0488) for several *DISC1* Interactome genes, but none survived correction for multiple testing (Supplementary Table S8).

#### Rare Functional Variant Analysis in the DISC1 Regulome

We analyzed the burden and accumulation rates of rare and singleton functional variants in the DISC1 Regulome. For SCZ compared to control samples, we observed a significantly increased burden of singleton disruptive variants (unadjusted *P*=0.0019, FWER_*within*_ *P*=0.0069, FWER_*across*_ *P*=0.0339, OR=1.3162, SE=0.0941; Figure 1 & Supplementary Table S9), and a nominally higher accumulation rate (4.13-fold, unadjusted *P*=9.00x10^−4^, FWER_*within*_ *P*=0.0185, FWER_*across*_ *P*=0.0965, Supplementary Table S10). In addition, the accumulation rate of rare disruptive variants, as opposed to singleton disruptive variants, was 3.47-fold higher in SCZ cases than in healthy controls and remained significant after multiple test correction (unadjusted *P*=1.68×10^−6^, FWER_*within*_ *P*=1.00×10^−4^, FWER_*across*_ *P*=0.0022, Supplementary Table S10). Although the burden of rare disruptive variants in SCZ was significant, and survived FWER correction for all tests within the trait, it did not meet the threshold for tests across all traits (unadjusted *P*=0.0061, FWER_*within*_ *P*=0.0228, FWER_*across*_ *P*=0.0863, Supplementary Table S9). We also observed a significantly higher proportion and burden of *NS*_*strict*_ singleton and rare variants in SCZ and disruptive singleton and rare variants in combined cases compared to controls, but none survived FWER for all tests across all traits (Supplementary Table S9 & S10). There was no evidence for an increased overall burden in rMDD, BD or combined cases compared to controls after FWER correction across all traits.

**Figure 1.**
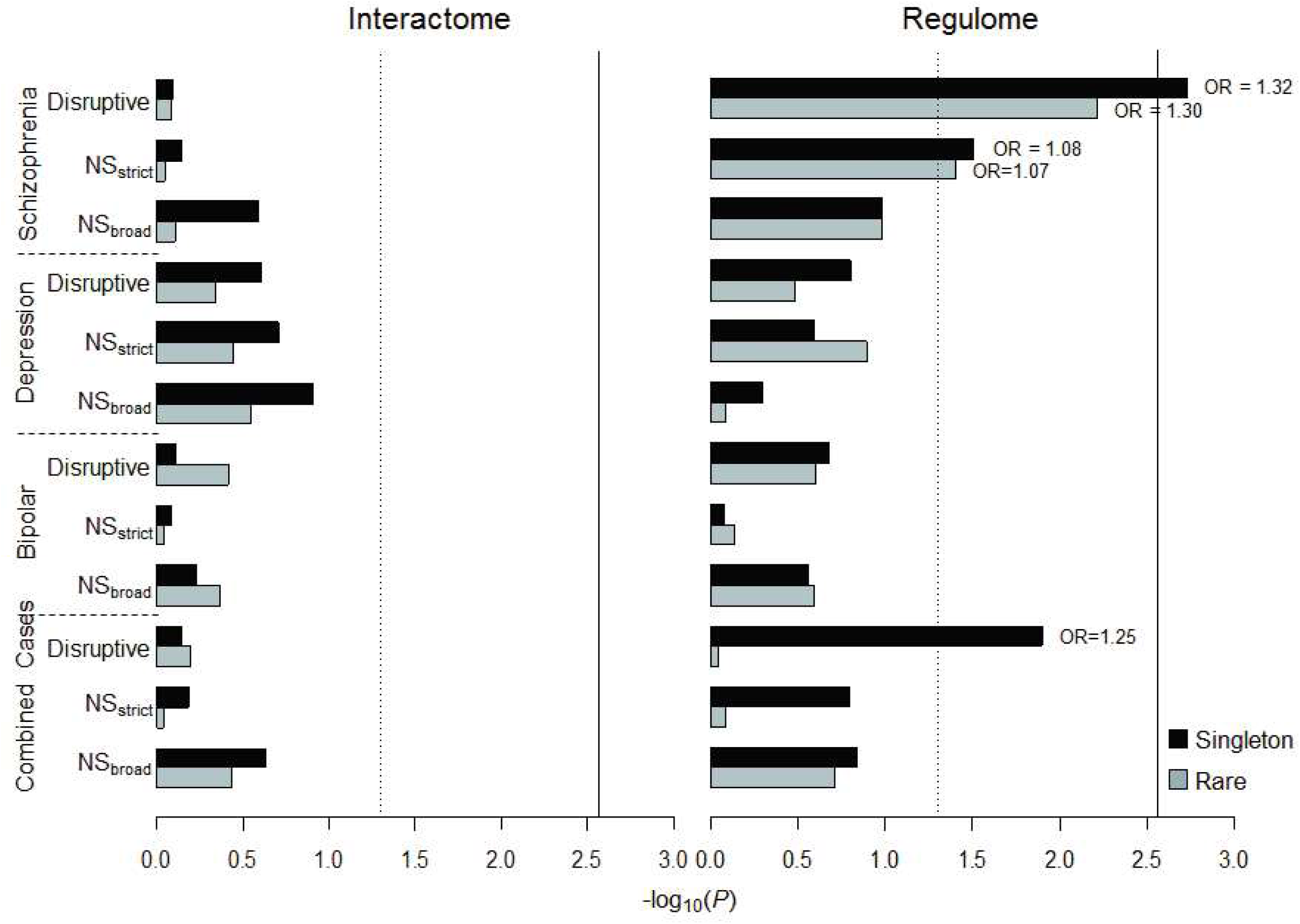
Gene set burden analysis of rare functional variants for case-control traits. Case-control gene set burden analysis of singletons and rare variants (MAF<1%) in the DISC1 Interactome (left) and Regulome (right). *x*-axis represents −log10(*P*), vertical dashed line: *P*=0.05, vertical solid line: FWER_*across*_ *P*=0.05; Odds ratio (OR) is labeled for the significant tests with *P*<0.05. Disruptive mutations, which included nonsense and splice site variants; NS_*strict*_, Non-synonymous strict damaging mutations which included disruptive variants plus missense variants predicted as damaging by all five algorithms (PolyPhen2 HumDiv and HumVar, SIFT, LRT and MutationTaster); NS_*broad*_, Non-synonymous broad damaging mutations which included disruptive plus missense variants predicted as damaging by at least one of the algorithms above.

At the gene-wide level, *Translin-associated factor X interacting protein 1 (TSNAXIP1)* showed greater burden of NS_*strict*_ singletons in rMDD (unadjusted *P*=1.29×10^−4^, FWER_*within*_ *P*=0.0253) and NS_*strict*_ rare variants in SCZ (unadjusted *P*=2.22×10^−4^, FWER_*within*_ *P*=0.0410, Supplementary Table S11) compared to controls. However this result did not survive correction for all tests (rMDD FWER_across_ *P*=0.0864, SCZ FWER_*across*_ *P*=0.1600). TSNAXIP1 has 16 exons encoding 712 amino acids. We validated 17 rare coding variants in *TSNAXIP1* in all carriers, including 1 splice site, 1 nonsense and 15 missense variants (Figure 2 & Supplementary Table S12). Of these 17 rare substitutions, 4 were previously reported in the 1000 Genomes Project European subset. In total, 7 rare variants in TSNAXIP1 including 2 disruptive and 5 predicted damaging missense variants contributed to the gene burden analysis of NS_*strict*_ variants in rMDD and SCZ. In a “leave-one-out” approach, we determined that the nonsense mutation at chr16:66405794 (rs146214814, R46X) located in exon 2, contributed most to the higher burden of NS_*strict*_ variants in SCZ. Relative to controls, this disruptive variant had a 3.58-fold higher allele frequency in SCZ (0.0146 vs 0.0041) and was not observed in any other mental illness cohort.

**Figure 2.**
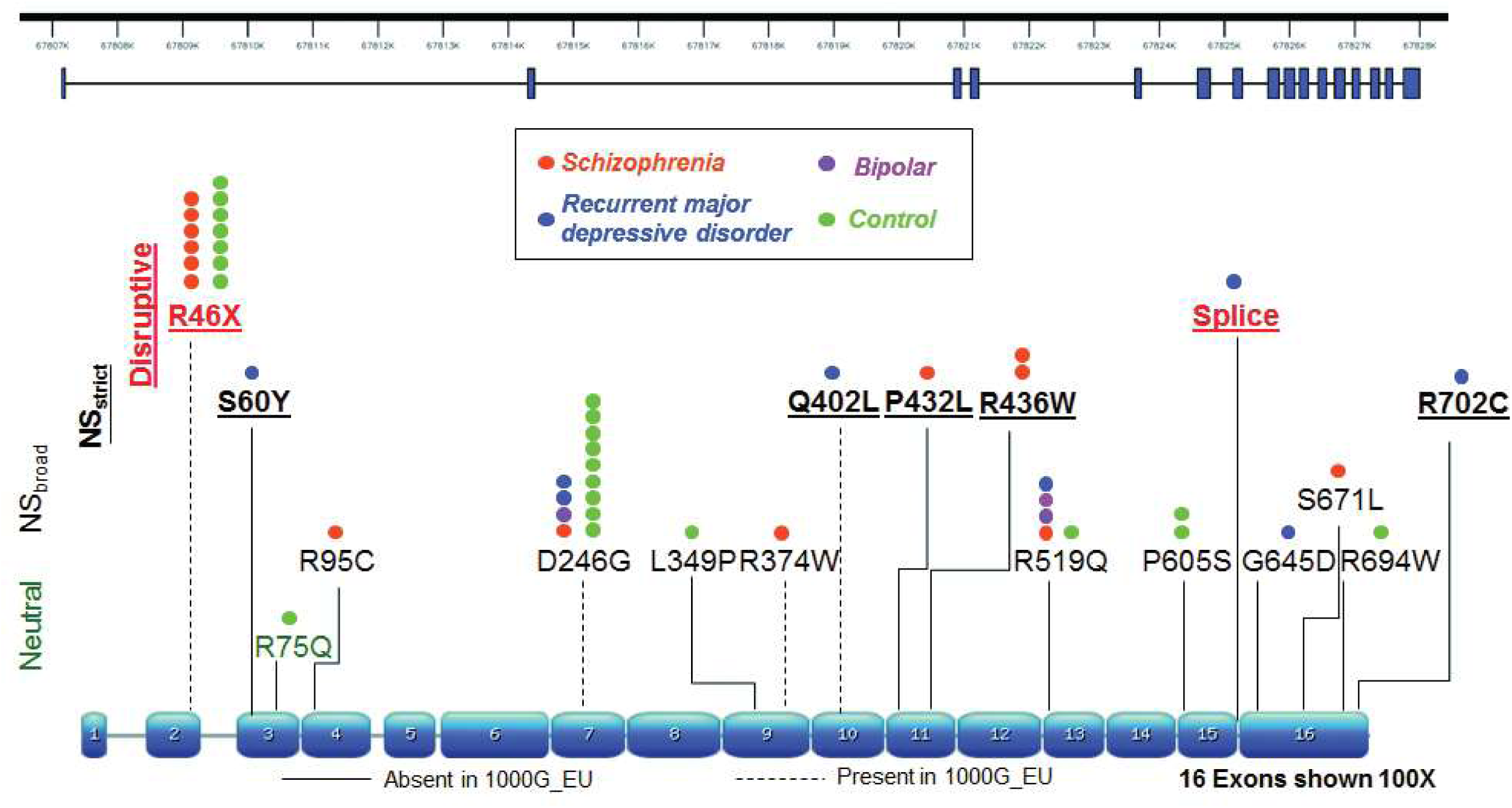
Translin-associated factor X interacting protein 1 *(TSNAXIP1)* rare functional variants. *TSNAXIP1* exon structure and mutations. Disruptive (red), nonsense and splice site variants; NS_*strict*_ (underlined), Non-synonymous strict damaging mutations were defined as disruptive plus missense variants predicted as damaging by all five algorithms (PolyPhen2 HumDiv and HumVar, SIFT, LRT and MutationTaster); NS_*broad*_, Non-synonymous broad damaging mutations were defined as disruptive plus missense variants predicted as damaging by at least one algorithm above. Neutral (green), neutral variants were defined as missense variants predicted as not damaging by all five algorithms above. Dash lines represent the variants present in the European subset of the 1000 Genomes Project (1000G_EU), and the solid lines represent the variants present in the 1000G_EU. The number of circles represents the number of samples carry the rare variant.

### Burden Analysis of Coding Variants on Quantitative Cognitive Ability and Personality Traits associated with Psychiatric Illness

We found that a significantly higher burden of singleton disruptive variants in the DISC1 Interactome was associated with lower cognitive ability assessed by Moray House Test (MHT) scores at age 11 (unadjusted *P*=9.35 × 10^−5^, FWER_*within*_ *P*=0.0005, FWER_*across*_ *P*=0.0043, *β*=−7.1141, SE=3.6863; Figure 3 & Supplementary Table S13). The burden of *NS*_*strict*_ singletons in the Interactome gene set was associated with lower MHT scores at age 11 (unadjusted *P*=0.0003, FWER_*within*_ *P*=0.0017, FWER_*across*_ *P*=0.0122, *β*=−2.7865, SE=1.2877). In addition, although these did not survive FWER_*across*_ correction, nominally significant associations in the burden of disruptive singletons were observed with MHT scores at age 70 (unadjusted *P*=0.0056, *β*=−6.6785), National Adult Reading Test (unadjusted *P*=0.0051, *β*=−6.9970) and General Fluid Intelligence (unadjusted *P*=0.0293, *β*=−0.5152). Interestingly, there were nominally significant associations between the burden of rare functional variants and increased symptoms of neuroticism (Disruptive singletons: unadjusted *P*=0.0154, *β*=6.5671), anxiety (*NS*_*strict*_ rare variants: unadjusted *P*=0.0349, *β*=0.1394) and depression (NS_*broad*_ singletons: unadjusted *P*=0.0431, *β*=0.2587). At the gene-wide level, no association was found between the variability in cognitive ability or personality scores and the burden of damaging or disruptive variants in any specific gene of the DISC1 Interactome after FWER_*across*_ correction (Supplementary Table S14).

**Figure 3.**
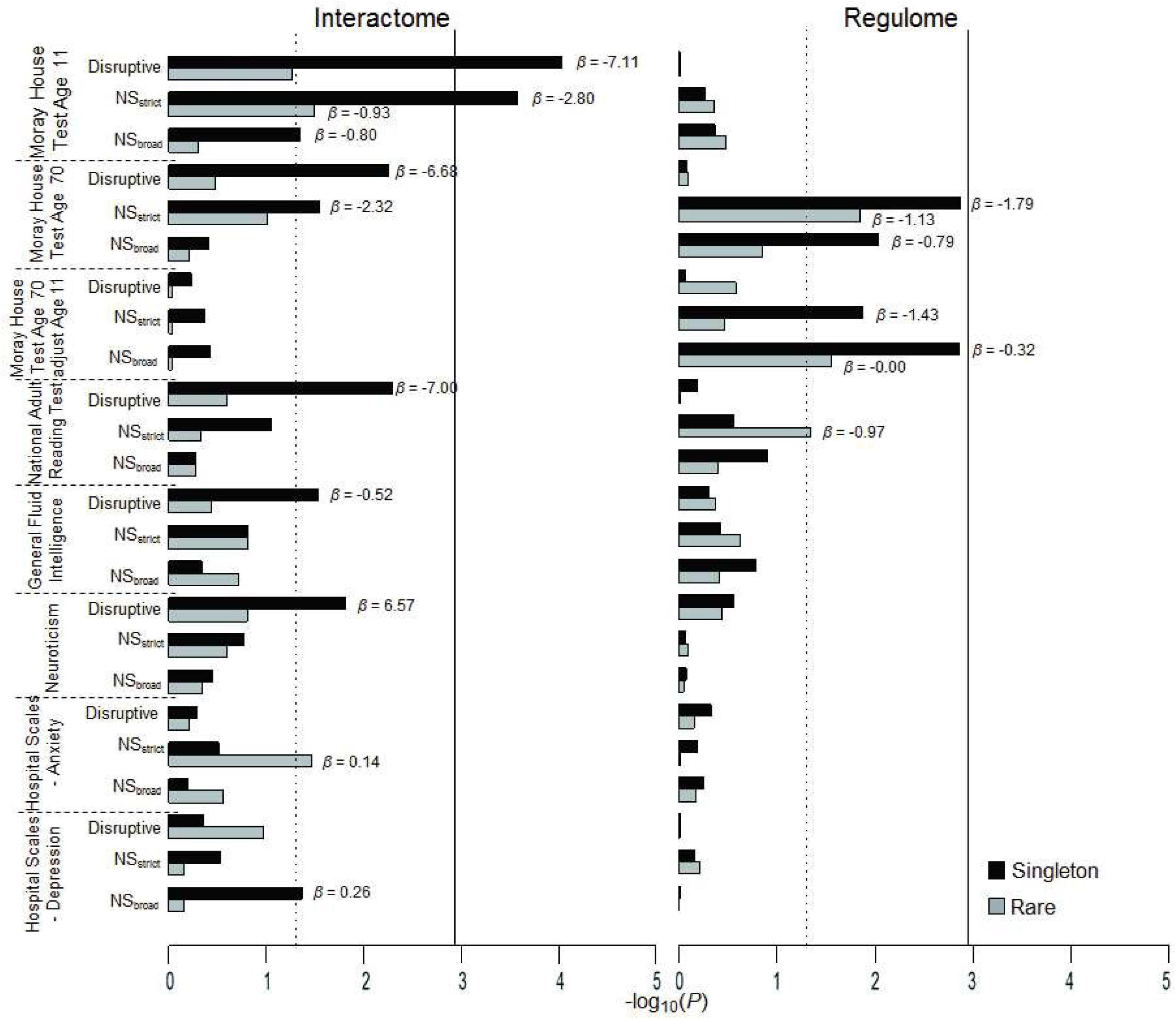
Gene set burden analysis of rare functional variants for quantitative traits. Quantitative trait gene set burden analysis of singletons and rare variants (MAF<1%) in the DISC1 Interactome (left) and Regulome (right). *x*-axis represents −log10(*P*), vertical dashed line: *P*=0.05, vertical solid line: FWER_*across*_ *P*=0.05; Effect size (*β*, Beta) is labeled for the significant tests with *P*<0.05. See phenotype descriptions in Supplementary Information for all quantitative traits. Moray House Test is the mental ability test used in the LBC1936 cohort. *A priori* hypothesis is that an increased burden of rare predicted damaging variants would reduce scores for cognitive variables and increase scores for personality traits. Disruptive, nonsense and splice site variants; NS_*strict*_, Non-synonymous strict damaging mutations were defined as disruptive plus missense variants predicted as damaging by all five algorithms (PolyPhen2 HumDiv and HumVar, SIFT, LRT and MutationTaster); NS_*broad*_, Non-synonymous broad damaging mutations were defined as disruptive plus missense variants predicted as damaging by at least one algorithm above.

In the analysis of the DISC1 Regulome, we observed a burden of NS_*strict*_ singletons associated with lower MHT scores at age 70 (unadjusted *P*=0.0014, *β*=−1.7895; Figure 3 & Supplementary Table S15) that withstood FWER correction for all tests within the trait (FWER_*within*_ *P*=0.0079), but not all tests across all traits (FWER_*across*_ *P*=0.0609). The burdens of NS_*strict*_ and NS_*broad*_ variants were nominally significantly associated with greater decrease in cognitive ability between the ages of 11 and 70 (NS_*strict*_ singletons: unadjusted *P*=0.0131, *β*=−1.4338; NS_*broad*_ singletons: unadjusted *P*=0.0014, *β*=−0.3175; NS_*broad*_ rare variants: unadjusted *P*=0.0280, *β*=−0.0010). At the gene-wide level, the strongest association with cognitive function was observed with rare and singleton NS_*strict*_ variants in *CACNA1C*, but this did not pass FWER_*across*_ correction (Supplementary Table S16).

## DISCUSSION

DISC1 has been a focus of attention since its first discovery, with interest growing as its impact on pathways and processes relating to psychopathology has been demonstrated experimentally.^24,26^ However, neither common variants in the *DISC1* gene, nor any of the DISC1 Interactome, are significantly associated with SCZ, BD or rMDD in large scale GWAS studies.^45,46^ The possible exception to this is the DISC1 interactor *PDE4B* that is the sole gene in a genome-wide significant region in the latest UK mega-analysis of schizophrenia in which.^47^ Rare variants of high impact provide a powerful complement to biological validation and insight. Recent case-control deep sequencing studies indicate that in individuals with SCZ rare loss-of-function variants are enriched in genes related to synaptic function,^30^ in target genes of the fragile X mental retardation protein^31^ and in genes known to be associated with SCZ.^48^ The biological impacts of several *DISC1* missense variants identified through deep sequencing have been demonstrated.^29,49^ We previously reported the discovery of rare disruptive *DISC1* variants in individuals with psychiatric illness and demonstrated the biological impact of the R37W variant.^17^ Here we report the association of both clinical diagnoses and cognitive ability with rare variants in the DISC1 gene and genes encoding known DISC1 interacting proteins, the DISC1 Interactome, and genes that show altered gene expression if the Interactome is disrupted, the DISC1 Regulome.

Before discussing these positive findings, we first consider some limitations of the study. Although the sample size was large by current standards, these numbers are modest in size for comprehensive rare variant detection.^17,50^ Burden analysis increases the power of analyses in such small samples, but the rules for annotating rare variants as ‘damaging’ are far from foolproof: biological validation is required. Last, but not least, whole genome sequencing of all 1 543 individuals, whilst ideal, was beyond the scope of our resources. Targeted capture sequencing was a practical option, but it is likely that relevant variants will have been missed by virtue of poor capture. It is also almost certainly the case that our list of *bona fide* DISC1 interactors is incomplete, and that contra wise, not all members of the Regulome that met our inclusion criteria will be regulated by DISC1 in practice.

Acknowledging these limitations, there were findings of note. No association was seen between rare variants in the DISC1 Interactome and any psychiatric diagnosis. There was, however, a significant excess of singleton disruptive variants in the DISC1 Regulome associated with SCZ, but not with BD or rMDD. We have shown that disruptive and NS_*strict*_ singleton variants in the DISC1 Interactome show significant association with cognitive ability at age 11. These classes of variants are also nominally associated in the DISC1 Regulome with cognitive ability at age 70 and change in cognitive ability between age 11 and 70. The DISC1 Regulome gene set was assembled from genes that show both i) altered expression in response to genetic variation in DISC1 or its interactors, and ii) evidence of association with psychiatric illness from candidate gene studies, or some of the earliest genome-wide association studies.^1,51,52^ It is therefore notable that we found association of rare Regulome variants with both increased schizophrenia risk and lower adult cognitive ability, particularly in older age. This mirrors the observation of association with common variants from the DISC1 Regulome in GWAS studies.^8,9,53,54^ Overall, the patterns of associations seen across diagnostic and cognitive traits in the DISC1 Interactome and Regulome are consistent with the hypothesis that genetic disruption of the DISC1 or its direct interactors has a proximal effect on cognitive ability and a distal effect, through regulation of gene expression, on schizophrenia risk in later life. Indeed, we have shown previously that disruption of the *Disc1* gene in mice results in altered expression of Nrxn1,^32^ a gene in which copy number variation, common variants and rare variants are associated with schizophrenia.^47,55,56^ The hypothesis of a distal effect of variants in the Interactome on disease risk is also consistent with the recent association of copy number variants linked to intellectual disability with schizophrenia in a much larger sample.^57^

The DISC1 Interactome is enriched for proteins involved in regulation of nervous system development, microtubule cytoskeleton organization, and vesicle localization (Supplementary Tables S17 & 18, Supplementary Figures S9-11). Our findings suggest that disruptive singletons in these biological processes may make significant contributions to variability in cognitive function. These processes have also been associated with intellectual disability.^58^ Together these datasets suggest that there is a spectrum of effect sizes or penetrance associated with genetic variants in this pathway. In contrast, the DISC1 Regulome is enriched for genes involved in synaptic transmission and glutamate-gated ion channel activity, reflecting the regulation by the DISC1 Interactome of these processes and their importance as inferred from GWAS. A role for DISC1 in glutamate-related processes has previously been suggested in both a mouse model and in the translocation family.^59,60^

In conclusion, and despite the limitations, these findings provide further genetic evidence to support the impact of both DISC1-interacting proteins and genes whose expression is modulated by genetic variants in the DISC1 pathway on psychopathology.

## CONFLICT OF INTEREST

WRM has participated in Illumina sponsored meetings over the past four years, and received travel reimbursement and an honorarium for presenting at these events. Illumina had no role in decisions relating to the study/work to be published, data collection and analysis of data and the decision to publish. WRM has participated in Pacific Biosciences sponsored meetings over the past three years and received travel reimbursement for presenting at these events. WRM is a founder and shareholder of Orion Genomics. WRM is a member of the scientific Advisory Board of RainDance, Inc.

## ACKNOWLEDGEMENTS

This study was supported by a gift from T and V Stanley and a grant from NIH (R01MH102068). We thank the LBC1936 participants and team members who contributed to these studies. Phenotype collection was supported by Age UK (The Disconnected Mind project). The work was undertaken by The University of Edinburgh Centre for Cognitive Ageing and Cognitive Epidemiology, part of the cross council Lifelong Health and Wellbeing Initiative (MR/K026992/1). Funding from the BBSRC and Medical Research Council (MRC) is gratefully acknowledged. ST thanks supports from the Howard University startup funds (U100193) and Junior Faculty Writing & Creative Works Summer Academy. ST acknowledges Professor Fatimah Jackson for critical comments.

## AUTHOR CONTRIBUTIONS

DJP and WRM designed the study, are PIs on the grant funding, supervised the study and supported the analysis. ST, SEM, PT, VM, MLC and MK carried out the bioinformatics analysis. MK, SM and ST organized and conducted targeted resequencing and variant validation. ST, PT, SEM, DJP and WRM wrote the manuscript, and all authors critically revised the manuscript.

